# Impact of Sugar-Based Baits on Midgut Microbiome Composition in *Aedes* Mosquitoes: Implications for Vector Control

**DOI:** 10.1101/2025.07.17.665286

**Authors:** Ranjitha Sambanthan, Nur Faeza Abu Kassim, Sara A. Abuelmaali, Wan Maryam Wan Ahmad Kamil, Sumiyyah Sabar, Kamarul Zaman Zarkasi, Wan Rosli Wan Ishak, Cameron E. Webb

**Affiliations:** 129 Medical Entomology Laboratory, School of Biological Sciences, Universiti Sains Malaysia, 11800 Minden, Penang, Malaysia; School of Physics, Universiti Sains Malaysia, 11800 Minden, Penang, Malaysia; Chemical Sciences Programme, School of Distance Education (SDE), Universiti Sains Malaysia, 11800 Minden, Penang, Malaysia; School of Health Sciences, Universiti Sains Malaysia, 16150 Kubang Kerian, Kelantan, Malaysia; Medical Entomology, NSW Health Pathology, Westmead Hospital, Westmead, NSW 2145, Australia; Sydney Institute for Infectious Diseases, University of Sydney, Westmead Institute for Medical Research, 176 Hawkesbury Road, Westmead, NSW 2145, Australia; BioG Expert Sdn Bhd, School of Biological Sciences, Universiti Sains Malaysia, 11800 Minden, Penang, Malaysia

**Keywords:** Bacteria, Culture, 16S rRNA gene, Midgut, Mosquitoes

## Abstract

Many severe mosquito-borne diseases are transmitted by the *Aedes sp.* mosquitoes. Control efforts have been strengthened by the implementation of novel integrated vector management techniques such as alginate hydrogel beads and appealing toxic sugar bait. These techniques help to control mosquitoes by taking advantage of their attraction to sugar. Different types of sugar that mosquitoes ingest during feeding can affect the makeup of microbiome in the midgut. Immune priming and baseline immune activity are maintained by mosquito midgut microbiome. Thus, this study is currently focusing on using microbial communities for vector control measures with a particular emphasis on how they consume various forms of sugar. Both wild and lab strain *Ae. aegypti* and *Ae. albopictus* mosquito samples were reared and fed with attractive targeted sugar baits (ATSBs) infused with *Chrysanthemum*, mango, mix and control solutions. Then, the impact on bacterial communities was assessed by using 16S rRNA gene sequences. According to our findings, the majority of the bacterial species in mango and mix treatments belonged to the Enterobacteriaceae family. A total of 24 various bacterial species were found in *Aedes* mosquitoes that fed on mango ATSBs. All isolates were members of three phyla from Actinobacteria (4.16%), Firmicutes (54.17%), and Proteobacteria (41.67%). Data reveals that different species, strains and diet affect the midgut bacterial diversity in the mosquitoes. In addition to improving our knowledge concerning the way this bacterium shapes the microbial community, a thorough investigation of the prevalence of the midgut bacterial community is essential for alerting present and future mosquito and disease control initiatives.

## 1.0 Introduction

The term “microbiome” refers to the grouping of all microorganisms that naturally reside on or within insect bodies including bacteria, fungi, viruses, and their genes. The microbiome is known to be a key interface between insect body and the environment. It will affect the current condition of the body by altering the response to environmental changes. While certain microorganisms operate as a buffer and reduce the toxicity of environmental contaminants and vice versa (National Institute of Environmental Health Sciences, 2025). The salivary glands, crop, gut, ovaries, Malpighian tubules, and hemocoel are among the many of the organs that mosquito microbial symbionts reside (Villegas et al., 2023). The developmental process and transmission of viruses and parasites rely on these organs. The gut is the main location of pathogen exposure due to the bloodmeal is storage and metabolization process in the mosquito’s midgut.

Gram-negative bacteria predominate in the gut microbiota of mosquitoes (Scolari et al., 2019). It was primarily composed of bacteria from the families Oxalobacteraceae, Enterobacteriaceae, Comamonadaceae, and genera Pseudomonas, Acinetobacter, Aeromonas, and Stenotrophomonas (David et al, 2016). According to reports, the midgut bacteria have a substantial impact on the capacity for disease transmission since they are linked to host-pathogen interaction and eventually, vector competency (Weiss et al., 2011). These results point to a unique disease control strategy by introducing particular bacterial strains with innate or designed anti-pathogen activity into the mosquito gut (Gusmão et al, 2010; Ramírez et al, 2012).

The identification and classification of Bacteria and Archaea communities is frequently accomplished by the use of 16S ribosomal RNA (rRNA) sequencing (Chun & Rainey, 2014). The 30S small subunit of a prokaryotic ribosome is made up of the bacterial 16S rRNA, which has a length of about 1500 bp (Fukuda et al., 2016). Generally, this gene consists of nine hypervariable regions interspersed with conserved sections. Several bacterial species share a high degree of conservation in the 16S rRNA gene’s conserved regions. The nine hypervariable regions (V1–V9) are informative for phylogenetic analysis and species identification because they show significant sequence diversity among various bacterial species and contain important information about genetic differences between bacterial species (Van de Peer et al., 1996).

The mosquito midgut represents the most important site for host-pathogen interaction that consist of the digestive tract which is responsible for their food digestion. The communities of microbiota play critical roles in their biological processes such as immune response function, sexual reproduction, digestion, development and refractoriness of pathogens (Huang et al., 2020). Thus, in this exploratory study, we aimed to expand the current understanding of the diversity of mosquito midgut microbiome before and after feeding on sugar sources.

## 2.0 Materials and methods

### 2.1 Samples collection and preparation

Both laboratory and wild strain of *Ae. aegypti* and *Ae. albopictus* were reared separately in different rearing room. Total of 15 male mosquitoes (three biological repeats) of laboratory strain *Ae. aegypti* were placed in different cages and fed with 30% w/v of mango, chrysanthemum, mix and control ATSBs in each cage respectively. The experiment was repeated for both male and female laboratory and wild strain *Ae. aegypti* and *Ae. albopictus*. The ATSBs were replaced every 24 hours because they were not treated with a preservative.

After three days, mosquitoes were collected using an aspirator and killed via cold shock, they were then transferred to sterile petri dishes for dissection. To ensure sterility, the dissection area was sanitized with 70% ethanol, using a dissecting microscope, the midguts of mosquitoes were carefully removed. For each species and sugar bait variant, five midguts from each replicate group and the control group were collected into 1.5 mL microcentrifuge tubes.

### 2.2 Isolation of microbial colonies from agar plate cultures

The collected midguts were sterilized with 300 µL of 70% ethanol for two minutes and rinsed three times with sterile distilled water. After removing the water, 100 µL of phosphate-buffered saline (PBS) was added to the tubes. The midguts were homogenized using a sterilized micropestle and the homogenates were serially diluted in PBS to seven-fold dilutions. The samples were streaked onto sterile nutrient agar media and incubated at 37°C for 24 to 48 hours under aseptic conditions. Plates without visible colonies were re-incubated for 16 hours to accommodate slow-growing bacteria. Colonies were classified by morphology size, shape, and color followed by subculture until pure colonies formed.

### 2.3 Bacterial Gram- staining

Gram staining process was carried out for identification of Gram-positive and Gram-negative bacterial colony. Bacterial smears were prepared by mixing a small quantity of colony with a drop of sterile distilled water on a clean glass slide. Slides were allowed to air dry before being heated under a Bunsen burner flame. The Gram staining procedure involved washing in distilled water in between each application of crystal violet (1 minute), iodine solution (1 minute), 70% ethanol for decolorization (5 seconds) and safranin (1 minute). After air drying, stained slides were observed at 100× magnification using an oil immersion light microscope. Gram-negative bacteria appeared pink or red, and Gram-positive bacteria were in purple (Beveridge, 2001).

### 2.4 DNA extraction

Approximately, 20mg of bacterial pellet was added in a 1.5ml of centrifuge tube accordingly for centrifuge at 8000 x g around 5 minutes to remove the supernatant. Then, lysis process was carried out differently for both Gram-positive and Gram-negative bacteria. For Gram-positive bacteria, the bacterial pellets were resuspended in 180μL bacteria pre-lysis buffer followed by addition of lysozyme with final concentration of 20mg/mL. The samples were incubated for 30 minutes at 37°C. Meanwhile, Gram-negative bacterial pellet was resuspended in 180μL TLB1 buffer.

The following steps are similar for Gram-positive and Gram-negative bacterial DNA extraction. Then, 25μL of Proteinase K solution was added to the tube and it were vortexed vigorously (Lin et al., 2021). Subsequently, all the tubes were sealed using parafilm and incubated for 3 hours at 56°C. The samples were vortexed every 15 minutes for the first two hours. After 3 hours, the samples were taken out from the incubator followed by addition of 200μL TLB2 buffer into it. The samples were then vortexed vigorously. Then, it was incubated at 70°C about 10 minutes. Afterwards, centrifugation was performed for the tubes contained insoluble particles at 11,000 x g. Then, the supernatant was transferred to a new 1.5mL centrifuge tube.

The next step is binding process. 210μL of 100% ethanol was added and mixed by vortex. Then, the lysate was transferred into a PrimeWay Genomic Column and was centrifuged at 11,000 x g for one minute. Subsequently, the PrimeWay Genomic Column was placed onto a new collection tube. The following step was washing the samples by adding 500μL Wash Buffer T1 into the column. Centrifugation was done with condition of 11,000 x g for one minute. The flow-through was discarded and the genomic column was placed back into collection tube. The same step was repeated by adding 600μL Wash Buffer T2 into it. For the drying process, the samples were centrifuge again for 1 minute to remove ethanol residue. Then, the genomic column was placed in a new centrifuge tube. Finally, 50μL of Elution Buffer added to the center of column membrane. The column was kept stand at room temperature for 1 minute and then followed by centrifugation to elute DNA.

### 2.5 PCR amplification and sequencing

All of the PCR components were assembled in a 0.2 mL PCR tube with PCR components of sterile distilled water (13.5 μL), DreamTaq Mastermix (12.5 μL), reverse primer (1 μL), forward primer (1 μL) and DNA template (2 μL). To produce a negative control, 2 μL of sterile distilled water was included to replace the DNA template. The 16S rRNA genes of their V3-V4 regions were amplified and sequenced on an Illumina HiSeq 2500 platform. The primer set used for this study were forward primer (27F), 5’ – GAG TTT GAT CCT GGC TCA – 3’ and reverse primer (1492R), 5’ – CGG CTA CCT TGT TAC GAC TT – 3’. PCR amplification was performed using T100 TM thermal cycler (Bio-Rad Laboratories, Inc, USA). The PCR cycling conditions that used were started with initial denaturation at 95 °C for 1 minute followed by denaturation at 95 °C for 30 seconds. Then, annealing process took place at 50 °C for 30 seconds and extension was at 72 °C for 1 minute. Denaturation, annealing and extension process were undergoes 35 cycles in total. Finally, the final extension process occurred at 72 °C for 5 minutes (Lee et al., 2019). The PCR products were analyzed by agarose gel electrophoresis.

### 2.6 Analysis of sequencing data

After the base calling analysis, the original data files from the sequencing platform were transformed into the original sequenced Reads Stored in FASTQ format. All of the sequences submitted to Genbank database. Sequences were analyzed using the Molecular Evolutionary Genetics Analysis (MEGA) software version 11.

## 3.0 Results

The results of bacterial species isolated from the midgut of female wild strain *Ae. aegypti* and *Ae. albopictus* mosquitoes under untreated and ATSBs treated conditions showed in Table 1. It shows that *Bacillus cereus* (PQ661268), *Bacillus albus* (PQ661254) and *Bacillus altitudinis* (PQ661256) dominated the microbiota of untreated wild female *Ae. aegypti*. On the other hand, female *Ae. albopictus* showed a unique microbial community that was mainly made up of *Microbacterium maritypicum* (PQ669681), *Acinetobacter nosocomialis* (PQ669684), and *Clostridium sporogenes* (PQ669689). This indicates a possibly pathogenic and more anaerobic bacterial composition. Both mosquito species were exposed to variety of gram-negative bacteria through the use of mango-based ATSBs. *Klebsiella pneumoniae* (PQ661365), *Klebsiella grimontii* (PQ661258) and *Klebsiella aerogenes* (PQ661262) are all the Enterobacteriaceae that renowned for their opportunistic pathogenicity and environmental adaptability, were shown to be abundant in the gut microbiota of *Ae. aegypti*. Bacteria such as *B. altitudinis* (PQ669661), *Stenotrophomonas maltophilia* (PQ670061) and *Micrococcus aloeverae* (PQ669216) colonized wild female *Ae. albopictus.* Interestingly, *B. altitudinis* was prevalent in both species, indicating that mango ATSBs were used for selective enrichment.

**Table 1.** Bacterial species isolated from the midgut of female wild strain *Ae. aegypti* and *Ae. albopictus* mosquitoes under untreated and ATSBs treated conditions.

|  | *Ae. aegypti* (wild strain female) |  | *Ae. albopictus* (wild strain female) |  |
| --- | --- | --- | --- | --- |
|  | Bacterial species | Accession number | Bacterial species | Accession number |
| Untreated | *Bacillus cereus* | PQ661268 | *Clostridium sporogenes* | PQ669689 |
|  | *Bacillus albus* | PQ661254 | *Acinetobacter nosocomialis* | PQ669684 |
|  | *Bacillus altitudinis* | PQ661256 | *Microbacterium maritipicum* | PQ669681 |
| Mango ATSBs | *Klebsiella pneumoniae* | PQ661365 | *Micrococcus aloeverae* | PQ669216 |
|  | *Klebsiella grimontii* | PQ661258 | *Stenotrophomonas maltophilia* | PQ670061 |
|  | *Klebsiella aerogenes* | PQ661262 | *Bacillus altitudinis* | PQ669661 |
| Chrysanthemum ATSBs | *Bacillus startosphaericus* | PQ661257 | *Delftia lacustris* | PQ669672 |
|  | *Bacillus velezensis* | PQ661260 | *Stenotrophomonas maltophilia* | PQ670839 |
|  | *Pseudomonas aeruginosa* | PQ661368 | *Stenotrophomonas pavanii* | PQ670082 |
| Mix ATSBs | *Enterobacter cloacae* | PQ661278 | *Paenibacillus* sp. | PQ669708 |
|  | *Pseudomonas protegens* | PQ661250 | *Bacillus cereus* | PQ671051 |
|  | *Pseudomonas aeruginosa* | PQ661269 | *Bacillus cereus* | PQ671077 |

Following exposure to *Chrysanthemum* ATSBs, *Pseudomonas aeruginosa* (PQ661368), *Bacillus startosphericus* (PQ661257) and *Bacillus velezensis* (PQ661260) dominated the gut profiles of wild female *Ae. aegypti*. *Delftia lacustris* (PQ669672), *S. maltophilia* (PQ670839) and *Stenotrophomonas pavanii* (PQ670082) were significantly enriched in the gut of *Ae. albopictus*. Meanwhile, *P. aeruginosa* (PQ661269), *Pseudomonas protegens* (PQ661250) and *E. cloacae* (PQ661278) were in the gut of *Ae. aegypti* treated with mixed ATSBs. The *B. cereus* (PQ671051, PQ671077) and *Paenibacillus sp.* (PQ669708) which are commonly found in soil and insect guts, were the two most prevalent bacterial taxa in wild female *Ae. albopictus*. Remarkably, *P. aeruginosa* appeared in both species, indicating that it continues to proliferate in mosquitoes treated with ATSBs.

Based on table 2, *Ae. aegypti* harbored microbiome consisting of *S. haemolyticus* (PQ676526), *B. amyloliquefaciens* (PQ678966) and *S. pasteuri* (PQ678886) in the untreated laboratory females. On the other hand, *M. maritypicum*, *K. aerogenes* (PQ681134), and *K. grimontii* (PQ681135) were found in *Ae. albopictus*. Notably, *M. maritypicum* was shared by both species, indicating overlap in the gut microbiota maintained in an untreated lab strain mosquito. Significant changes in microbiota have been driven about by the sugar baits made from mangoes. Along with *P. aeruginosa* (PQ681124), *B. cereus* (PQ678965), *B. tropicus* (PQ678967) and *B. subtilis* (PQ678969) were the predominant bacteria in *Ae. aegypti*. The *C. botulinum* (PQ681125), *S. maltophilia* (PQ681126) and *P. aeruginosa* were all present in *Ae. albopictus*. The two species shared *P. aeruginosa*, suggesting that mango ATSBs facilitate environment for gram-negative bacteria to colonize.

**Table 2.** Bacterial species isolated from the midgut of female lab strain *Ae. aegypti* and *Ae. albopictus* mosquitoes under untreated and ATSBs treated conditions.

|  | *Ae. aegypti* (lab strain female) |  | *Ae. albopictus* (lab strain female) |  |
| --- | --- | --- | --- | --- |
|  | Bacterial species | Accession number | Bacterial species | Accession number |
| Untreated | *Staphylococcus haemolyticus* | PQ676526 | *Microbacterium maritipicum* | PQ681133 |
|  | *Bacillus amyloliquefaciens* | PQ678966 | *Klebsiella aerogenes* | PQ681134 |
|  | *Staphylococcus pasteurii* | PQ678886 | *Klebsiella grimontii* | PQ681135 |
| Mango ATSBs | *Bacillus cereus* | PQ678965 | *Pseudomonas aeruginosa* | PQ681124 |
|  | *Bacillus tropicus* | PQ678967 | *Clostridium botulinum* | PQ681125 |
|  | *Bacillus subtilis* | PQ678969 | *Stenotrophomonas maltophilia* | PQ681126 |
| Chrysanthemum ATSBs | *Lactobacillus iners* | PQ678970 | *Chryseobacterium indologens* | PQ681127 |
|  | *Bacillus paramycoides* | PQ678885 | *Chryseobacterium aureum* | PQ681128 |
|  | *Chryseobacterium indologens* | PQ675785 | *Bacillus velezensis* | PQ681129 |
| Mix ATSBs | *Enterobacter asburiae* | PQ675780 | *Klebsiella grimontii* | PQ681130 |
|  | *Aeromonas veronii* | PQ675812 | *Aeromonas veronii* | PQ681131 |
|  | *Bacillus cereus* | PQ678956 | *Klebsiella pneumoniae* | PQ681132 |

*Ae. aegypti* was a vector of *L. iners* (PQ678970), *B. paramycoides* (PQ678885) and *C. indologenes* (PQ675785) in the *Chrysanthemum* group. However, *B. velezensis* (PQ681129), *C. aureum* (PQ681128) and *C. indologens* (PQ681127) were linked to *Ae. albopictus.* After being exposed to *Chrysanthemum* ATSBs, both mosquito species showed the presence of *Chryseobacterium spp*., indicating that floral ATSBs play a major role in the acquisition of plant-associated bacteria. The *Ae. aegypti* mosquitoes treated with mixed ATSBs showed indications of *B. cereus* (PQ678956), *Aeromonas veronii* (PQ675812) and *E. asburiae* (PQ675780). The *A. veronii* (PQ681131), *K. grimontii* (PQ681130) and *K. pneumoniae* (PQ681132) were all present in *Ae. albopictus*. The ability of mixed floral baits to transfer water-associated opportunistic bacteria into the gut is suggested by the consistent finding of *A. veronii* in both species.

Table 3 shows that the bacterial population in the wild strain male *Ae. aegypti* untreated group was primarily made up of *B. cereus* (PQ661266, PQ661267) and *B. tropicus* (PQ661264). On the other hand, *L. komagatae* (PQ671410), *B. tropicus* (PQ671280) and *B. cereus* (PQ671709) were found in *Ae. albopictus*. Interestingly, both mosquito species shared *B. cereus* and *B. tropicus*, indicating that these bacterial taxa maybe a natural component of the microbiota in male wild populations. Bacteria such as *E. cloacae* (PQ661157), *B. amyloliquefaciens* (PQ661377) and *B. cereus* (PQ661373) colonized *Ae. aegypti* treated with mango ATSBs. Similarly, *E. coli* (PQ668632) and *E. cloacae* (PQ668541, PQ670976) were repeatedly found in *Ae. albopictus*. Given that *E. cloacae* is frequently found in both species and it is likely due to the sugar-rich formulation of mango ATSBs favors facultative anaerobes, which in turn promotes the growth or acquisition of enteric bacteria.

**Table 3.** Bacterial species isolated from the midgut of male wild strain *Ae. aegypti* and *Ae. albopictus* mosquitoes under untreated and ATSBs treated conditions.

|  | *Ae. aegypti* (wild strain male) |  | *Ae. albopictus* (wild strain male) |  |
| --- | --- | --- | --- | --- |
|  | Bacterial species | Accession number | Bacterial species | Accession number |
| Untreated | *Bacillus cereus* | PQ661267 | *Leucobacter komagatae* | PQ671410 |
|  | *Bacillus cereus* | PQ661266 | *Bacillus tropicus* | PQ671280 |
|  | *Bacillus tropicus* | PQ661264 | *Bacillus cereus* | PQ671709 |
| Mango ATSBs | *Enterobacter cloacae* | PQ661157 | *Enterobacter cloacae* | PQ668541 |
|  | *Bacillus amyloliquefaciens* | PQ661377 | *Enterobacter cloacae* | PQ670976 |
|  | *Bacillus cereus* | PQ661373 | *Escherichia coli* | PQ668632 |
| Chrysanthemum ATSBs | *Bacillus licheniformis* | PQ661367 | *Chryseobacterium bernardetii* | PQ668615 |
|  | *Microbacterium paraoxydans* | PQ661363 | *Chryseobacterium indologenes* | PQ668617 |
|  | *Bacillus thuringiensis* | PQ661366 | *Chryseobacterium rhizoplanae* | PQ668633 |
| Mix ATSBs | *Bacillus thuringiensis* | PQ661372 | *Aerococcus viridans* | PQ670297 |
|  | *Bacillus paramycoides* | PQ661364 | *Enterobacter ludwigii* | PQ669186 |
|  | *Bacillus thuringiensis* | PQ661378 | *Pantoea dispersa* | PQ671643 |

*B. licheniformis* (PQ661367), *M. paraoxydans* (PQ661363) and *B. thuringiensis* (PQ661366) were among the microbes that present in *Ae. aegypti* exposed to *Chrysanthemum* ATSBs. In contrast, *C. bernardetii* (PQ668615), *C. indologenes* (PQ668617) and *C. rhizoplanae* (PQ668633) were linked to *Ae. albopictus*. This tendency suggests a divergence in bacterial uptake between species, as wild male *Ae. albopictus* prefers *Chryseobacterium spp*., which are known to be prevalent in plant-associated settings. The microbiota of *Ae. aegypti* mosquitoes treated with mixed ATSBs including *A. viridans* (PQ670297), *B. thuringiensis* (PQ661372, PQ661378) and *B. paramycoides* (PQ661364). Meanwhile, *P. dispersa* (PQ671643) and *E. ludwigii* (PQ669186) were found in *Ae. albopictus*. Notably, enteric bacteria like *Pantoea* and *Enterobacter* predominated in *Ae. albopictus*, but *B. thuringiensis* was frequently found in wild male *Ae. aegypti*.

According to table 4, three different bacterial species were found in untreated male *Ae. aegypti* laboratory mosquitoes such as *B. cereus* (PQ678982), *M. endophyticus* (PQ678981) and *B. thuringiensis* (PQ678980). In contrast, *A. variabilis* (PQ681121), *B. albus* (PQ681122) and *B. tropicus* (PQ681123) dominated the bacterial profile of lab strain male *Ae. albopictus*. Under laboratory circumstances, there was no evidence of species overlap between the two species in the untreated group, suggesting species-specific core microbiota. Microbial profiles of the two species showed both overlap and divergence as a result of exposure to mango ATSBs. Three bacterial species were identified in *Ae. aegypti* consist of *B. sonorensis* (PQ678973), *C. botulinum* (PQ678972) and *B. subtilis* (PQ678971). Nevertheless, *B. thuringiensis* (PQ681114) and *B. cereus* (PQ681112, PQ681113) dominated lab strain male *Ae. albopictus*. Interestingly, *B. cereus* was present in both mosquito species, indicating that the sugar bait media cause impact on bacterial colonization.

**Table 4.** Bacterial species isolated from the midgut of male lab strain *Ae. aegypti* and *Ae. albopictus* mosquitoes under untreated and ATSBs treated conditions.

|  | *Ae. aegypti* (lab strain male) |  | *Ae. albopictus* (lab strain male) |  |
| --- | --- | --- | --- | --- |
|  | Bacterial species | Accession number | Bacterial species | Accession number |
| Untreated | *Bacillus thuringiensis* | PQ678980 | *Acinetobacter variabilis* | PQ681121 |
|  | *Micrococcus endophyticus* | PQ678981 | *Bacillus albus* | PQ681122 |
|  | *Bacillus cereus* | PQ678982 | *Bacillus tropicus* | PQ681123 |
| Mango ATSBs | *Bacillus subtilis* | PQ678971 | *Bacillus cereus* | PQ681112 |
|  | *Clostridium botulinum* | PQ678972 | *Bacillus cereus* | PQ681113 |
|  | *Bacillus sonorensis* | PQ678973 | *Bacillus thuringiensis* | PQ681114 |
| Chrysanthemum ATSBs | *Bacillus cereus* | PQ678974 | *Chryseobacterium aureum* | PQ681115 |
|  | *Micrococcus luteus* | PQ678975 | *Bacillus subtilis* | PQ681116 |
|  | *Bacillus cereus* | PQ678976 | *Chryseobacterium indologenes* | PQ681117 |
| Mix ATSBs | *Bacillus cereus* | PQ678977 | *Lysinibacillus fusiformis* | PQ681118 |
|  | *Bacillus cereus* | PQ678978 | *Staphylococcus epidermidis* | PQ681119 |
|  | *Bacillus cereus* | PQ678979 | *Staphylococcus warneri* | PQ681120 |

After being exposed to *Chrysanthemum* ATSBs, *Ae. aegypti* showed relatively diversified microbiota including *C. aureum* (PQ681115), *B. cereus* (PQ678974, PQ678976) and *M. luteus* (PQ678975). The bacteria *B. subtilis* (PQ681116), *C. indologenes* (PQ681117) and *C. aureum* (PQ681115) were among the microbiota in *Ae. albopictus* midgut. The multiple occurrence of *C. aureum* indicates that some bacteria originated from flowers may be able to colonize different mosquito species. Significant taxonomic overlap was seen in the microbiome of both mosquito species exposed to mixed ATSBs. *B. cereus* (PQ678977, PQ678978, and PQ678979) and *Lysinibacillus fusiformis* (PQ681118) were repeatedly isolated from *Ae. aegypti*. Along with *B. cereus*, *Ae. albopictus* also shows the presence of *S. epidermidis* (PQ681119) and *S. warneri* (PQ681120). The high prevalence of *B. cereus* in both species highlights its ecological competitiveness and resistance in sugar-rich habitats.

A total of 24 various bacterial species were found in *Aedes* mosquitoes that fed on mango ATSBs (Figure 1). All isolates were members of three phyla from Actinobacteria (4.16%), Firmicutes (54.17%), and Proteobacteria (41.67%). *Micrococcus aloeverae* from phyla Actinobacteria only present in female ALW. Bacteria from wild strain female *Ae. albopictus* were from three different phylum such as Proteobacteria (*Stenotrophomonas maltophilia*), Firmicutes (*Bacillus altitudinis*) and Actinobacteria (*Micrococcus aloeverae*). The midgut of mango ATSBs fed laboratory strain male and female *Ae. aegypti* and male *Ae. albopictus* fully colonized by phyla Firmicutes. Genus *Bacillus* dominated the midgut of these mosquitoes followed by genus *Clostridium*. However, phyla Proteobacteria fully occupied the midgut of wild strain female *Ae. aegypti* and male *Ae. albopictus*. The dominant bacterial species identified were *Bacillus cereus* (16.67%) followed by *Enterobacter cloacae* (12.5%).

**Figure 1.**
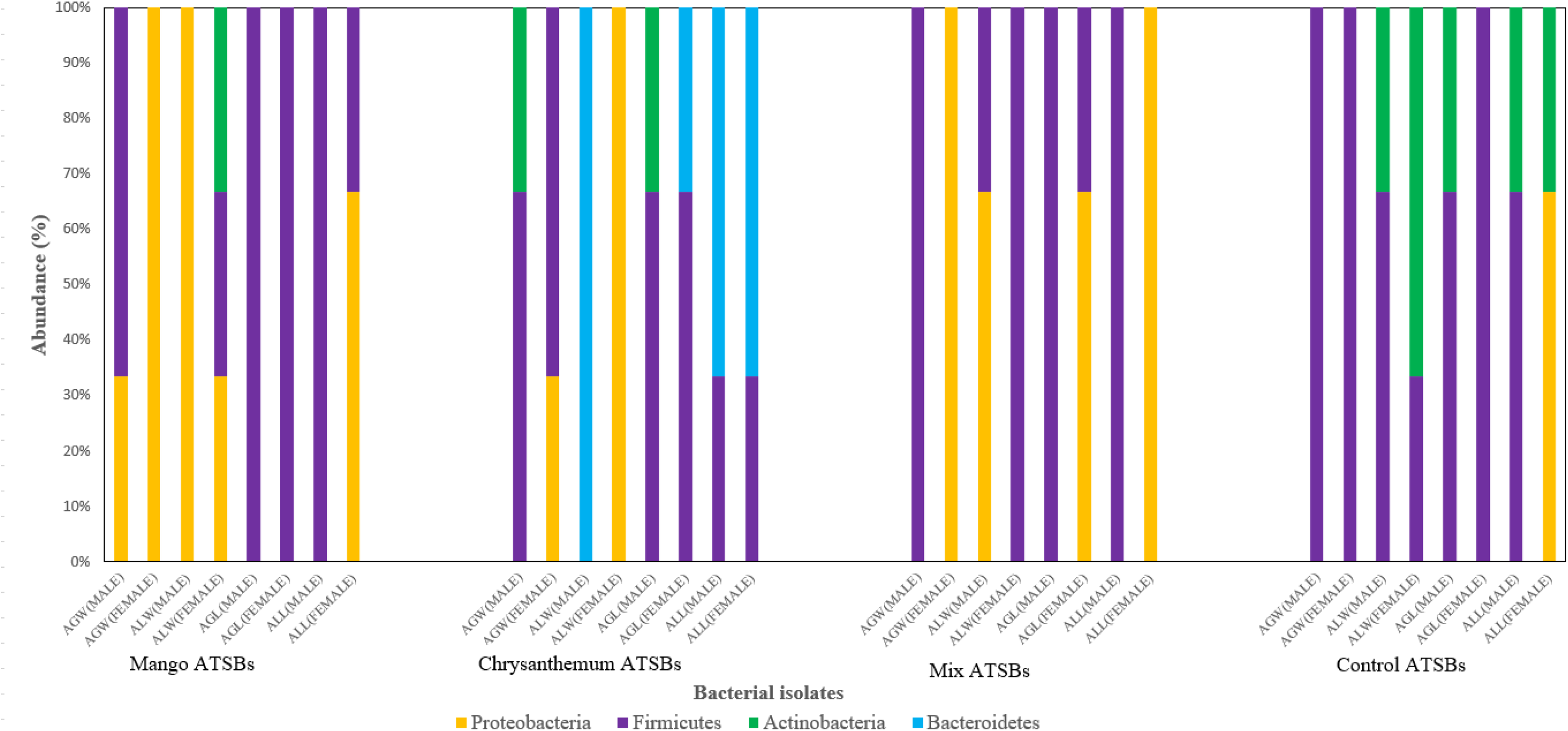
Abundance of identified bacterial species exposed to different ATSBs classified according to their respective phylum isolated from midgut of wild and laboratory strain *Ae. aegypti* and *Ae. albopictus*.

According to Figure 1, bacterial species isolated from midgut of *Aedes* mosquitoes that fed on *Chrysanthemum* ATSBs were from four different phylum. All isolates were members of Actinobacteria (8.33%), Firmicutes (41.67%), Bacteroidetes (33.33%) and Proteobacteria (16.67%). All of the bacteria from phylum Firmicutes are under genus *Bacillus* (100%) while genus *Chryseobacterium* (100%) dominates phylum Bacteroidetes. The data shows that female ALW consist of bacteria from phylum Proteobacteria such as *Delftia lacustris*, *Stenotrophomonas maltophilia* and *Stenotrophomonas* pavanii. However, male ALW only contains microbiome from phylum Bacteroidetes such as *C. bernadetii*, *C. indologenes* and *C. rhizoplanae*.

Furthermore, the microbiome isolated from the midgut of *Aedes* mosquitoes after exposed to mix ATSBs indicate that there were only two phylum presence which are Firmicutes (58.33%) and Proteobacteria (41.67%). Phylum Firmicutes consist of genus *Bacillus*, *Clostridium* and *Staphylococcus* meanwhile phylum Proteobacteria consist of genus *Aeromonas*, *Pantoea*, *Enterobacter*, *Pseudomonas* and *Klebsiella*. The midgut of female AGW and ALL completely colonized by phylum Proteobacteria. In addition, male AGW, AGL and ALL including female ALW predominantly established by phylum Firmicutes. Among all of the microbiome, *Bacillus cereus* have the highest abundance 20.83% in total. There are other *Bacillus* species found in the midgut of *Aedes* mosquitoes such as *Bacillus paramycoides* and *Bacillus thuringiensis*.

The mosquitoes fed on control ATSBs confirm the presence of three different phylum namely Proteobacteria (8.33%), Actinobacteria (25%) and Firmicutes (66.67%). Male AGW and AGL together with female AGL only indicate the Firmicutes group bacteria (100%) in the midgut of the mosquitoes. Phylum Proteobacteria only presence in female ALL which is represented by *Klebsiella grimontii* and *Pseudomonas aeruginosa*. Example of Phylum Actinobacteria that found in the mosquito midgut are *Microbacterium maritypicum*, *Micrococcus endophyticus*, and *Leucobacter komagatae*. Genus Bacillus and Staphylococcus from phylum Firmicutes are the dominant bacterial species species group in the control ATSBs fed mosquito midgut.

## 4.0 Discussion

The *Aedes* mosquitoes are known to be a major vector of arboviral illnesses, which raises serious issues for public health. Therefore, their midgut microbiome communities especially those related to sugar variants are very important for recognizing the function of microbiota in vector-parasite interactions. A clear difference between the identified bacterial species before and after treatment were highlighted in this study.

The microbial variation may impact vector competence, survivability, physiology and the ecological fitness of mosquito populations. According to our findings, the majority of the bacterial species in the mango and mix treatments belonged to the Enterobacteriaceae family. *Klebsiella* and *Enterobacter* had the highest number of bacterial genera. The bacterial species included *Klebsiella pneumoniae*, *Klebsiella grimontii*, *Enterobacter ludwigii*, *Enterobacter cloacae* and *Enterobacter asburiae*. Prior research has repeatedly shown that the most common bacteria isolated from *Aedes* mosquitoes were *Enterobacter* species (Lin et al., 2021; Yadav et al., 2015). The recent discovery of *E. cloacae* is significant since multiple investigations have shown they prevent *Plasmodium* from developing in the midgut of *Anopheles* mosquitoes (Ezemuoka et al., 2020; Cirimotich et al., 2011). *Enterobacter cloacae*, a common gut symbiont in mosquitoes, plays a crucial role in modulating the mosquito’s physiology and vector competence. It colonizes the mosquito midgut and contributes to the maintenance of gut homeostasis by aiding in nutrient metabolism and stimulating immune responses. Notably, *E. cloacae* can inhibit the development of pathogens such as *Plasmodium* spp., the causative agents of malaria, by inducing the mosquito’s innate immune pathways and generating reactive oxygen species that are hostile to the parasites. This natural antagonism has made *E. cloacae* a candidate for paratransgenic strategies, where it is genetically engineered to express anti-pathogen effector molecules, thereby reducing the mosquito’s ability to transmit diseases (Cirimotich et al., 2011). Furthermore, it was found that *E. cloacae* can be used to control Leishmania in the sand fly *Phlebotomus papatasito* through the method of paratransgenesis (Maleki-Ravasan et al., 2015). In addition, it has been demonstrated that *E. cloacae* may effectively transfer, express, and spread foreign genes within termite colonies (Husseneder & Grace, 2005). These results suggest that using *E. cloacae* directly or via paratransgenic methods is possible for controlling vector-borne illnesses and lowering the spread of pathogens.

Since *Klebsiella* is a frequent genus of bacteria identified in the midgut of *Aedes* mosquitoes, its discovery is also very crucial. *Klebsiella* spp., particularly *K. pneumoniae* and *K. oxytoca* are common gut symbionts in mosquitoes and play roles in host development, immunity, and pathogen interference. The symbiotic interaction between *Ae. aegypti* mosquitoes and *Klebsiella spp.* was examined in a prior study (Mosquera et al., 2021). The results showed that volatile organic compounds (VOCs) released by *Klebsiella* species significantly influence the mosquitoes’ choices on oviposition. These VOCs had a significant impact on the gravid mosquitoes to choose their breeding grounds, which may have an impact on the dynamics of their population and the spread of diseases carried by mosquitoes. It is possible to create attract and kill mosquito management system by incorporate with VOCs which promotes mosquito oviposition.

These bacteria aid in nutrient metabolism and nitrogen fixation, which supports mosquito larval growth and adult fecundity. Additionally, they can stimulate the mosquito immune system, leading to enhanced production of antimicrobial peptides and reactive oxygen species, which are detrimental to invading pathogens such as *Plasmodium* and arboviruses. Due to these properties and their ability to stably colonize the mosquito gut, *Klebsiella* spp. have also been investigated as paratransgenic agents for vector control, engineered to deliver anti-pathogen molecules within the mosquito midgut (Gusmão et al., 2010; Tchioffo et al., 2016).

According to Kyritsis et al., (2017), *Klebsiella pneumoniae* in this genus which has been used as a biotechnological approach for controlling and sterilizing insects particularly to manage pests such as the Mediterranean fruit fly (Kyritsis et al., 2017). In grasshoppers, this bacteria species has also been found to be endosymbiotic (Shi et al., 2011). Furthermore, *Klebsiella variicola* has been often isolated from a variety of human sources and is known to be a pathogenic species for humans. In fruit crops, it has also been reported to be an endosymbiont of insects (Barrios-Camacho et al., 2019).

Numerous genera including Enterobacter, *Klebsiella*, *Pantoea*, *Acinetobacter*, *Pseudomonas*, *Bacillus*, *Staphylococcus*, *Micrococcus* and *Aeromonas* have been frequently isolated from the midgut of various mosquito species among the bacterial isolates found in this study (Moro et al., 2013; Boissière et al., 2012 & Zouache et al., 2011). In particular, the midgut of *Ae. aegypti* and *Ae. albopictus* collected from a geographically remote location in the past has also yielded the following bacteria. For instance, *Aeromonas veronii*, *Bacillus aerophilus*, *Bacillus cereus*, *Enterobacter asburiae*, *Enterobacter cloacae*, *Klebsiella pneumoniae*, *Pantoea dispersa*, *Pseudomonas aeruginosa*, and *Stenotrophomonas maltophilia* (Yadav et al., 2015). This indicating that numerous microbiomes established themselves by becoming commensal and play significant roles in the life cycle of vectors (Chandel et al., 2013).

Despite other factors, it has been demonstrated that the genders of the mosquitoes are connected to the variety of the midgut microbiota (Moro et al., 2013). This fact is in line with this study. Several bacterial species were solely found in the female mosquitoes such as *Micrococcus aloeverae*, *Staphylococcus haemolyticus*, *Staphylococcus pasteuri*, *Aeromonas veronii* and *Microbacterium maritypicum*. However, some bacterial species were exclusively found in male mosquitoes such as *Clostridium botulinum*, *Micrococcus endophyticus*, *Staphylococcus warneri*, *Lysisnibacillus fusiformis* and *Acinetobacter variabilis*. Microbial diversity was consistently higher in female mosquitoes across both species and strains, corroborating earlier findings that link greater bacterial richness with hematophagous behavior (Dennison et al., 2014). Female mosquitoes are exposed to a wider array of microbial sources through blood feeding and oviposition behaviours, which may facilitate the acquisition of gut-associated opportunistic bacteria such as *Clostridium*, *Chryseobacterium*, and *Acinetobacter* spp. Conversely, male mosquitoes that reliant on sugar meals exhibited a gut microbiome primarily dominated by *Bacillus* and other Firmicutes, likely reflecting restricted dietary intake and fewer environmental interactions (Apte-Deshpande et al., 2012).

A pronounced divergence in microbial diversity was observed between laboratory and wild strain mosquitoes. Both male and female wild strain mosquitoes harboured a broader range of taxa including *Paenibacillus*, *Aeromonas*, *Enterobacter*, and *Acinetobacter* spp. These genera are associated with natural habitats and organic material, suggesting environmental exposure as a key driver of microbial diversity (Minard et al., 2013). *Paenibacillus* is known to produce antimicrobial peptides and fix nitrogen, which may aid mosquitoes in maintaining nutritional homeostasis (Dantur et al., 2015). Laboratory strains showed reduced diversity, with a microbial profile largely dominated by Firmicutes. This reduced complexity may be attributed to sterile rearing conditions, consistent artificial diets, and a lack of environmental microbial reservoirs (Coon et al., 2014).

Across all conditions, members of the genus *Bacillus*, particularly *B. cereus*, *B. thuringiensis*, and *B. subtilis*, were frequently isolated from both mosquito species and sexes. The dominance of *Bacillus* spp. aligns with previous studies that identified these spore-forming Firmicutes as common gut symbionts, likely due to their environmental robustness and metabolic versatility (Ruiu, 2015). Benefits of these bacteria including increased tolerance to environmental stress and better production of digestive enzymes in the gut (Gusmão et al., 2010). Meanwhile, the role of *Bacillus thuringiensis* is recorded for the synthesis of toxins (Sarrafzadeh et al., 2005).

*Klebsiella* species such as *K. pneumoniae*, *K. grimontii*, *K. aerogenes* were prevalent in female mosquitoes, especially those exposed to mango ATSBs, suggesting potential floral or environmental acquisition. Notably, *Pseudomonas aeruginosa* and *Stenotrophomonas maltophilia* were frequently identified in female wild strains and post ATSBs exposure, possibly reflecting adaptive colonization of sugar rich gut environments or residual microbial presence from plant baits (Patil et al., 2022).

The introduction of floral-based sugar baits significantly altered the gut microbiota of mosquitoes. Mango ATSBs promoted colonization by facultative anaerobes such as *Klebsiella*, *Clostridium*, and *Enterobacter* organisms that known for their metabolic adaptability and capacity to utilize diverse carbohydrate sources (Benedek et al., 2016). Chrysanthemum ATSBs increased the abundance of *Chryseobacterium*, *Stenotrophomonas*, and *Pseudomonas* spp., bacteria which associated with environmental and plant microbiota that contribute antimicrobial metabolites (Ruiu, 2015). Mixed ATSBs introduced the broadest spectrum of bacteria including *Paenibacillus*, *Aeromonas*, and *Pantoea* spp., suggesting that compound floral blends may enhance microbial colonization through nutritional diversity or synergistic bacterial interactions.

In addition to the bacterial species listed above, we isolated *Chryseobacterium rhizoplanae, Microbacterium paraoxydans*, and *Clostridium sporogenes* from the midgut of mosquitoes during the investigation. Kämpfer and colleagues identified *C. rhizoplanae* a Gram-negative bacterium species, from the rhizoplane of maize after isolating it from current blood-fed individuals (Kämpfer et al., 2015). The *M. paraoxydans* which was isolated in this investigation was first identified in a Belgian patient with acute lymphoblastic leukemia and it is also identified in clinical samples and fish (Laffineur et al., 2003; Soto-Rodriguez et al., 2013). The *Clostridium sporogenes* species was first identified from human feces was later reported from the gastrointestinal system of humans and other mammals (Oakeson et al., 2016).

## 5.0 Conclusion

In summary, our research concentrated on the microbiome communities found in both wild and laboratory strain *Aedes* mosquito midgut and specifically examining their activity towards various sugar. We examined the effects of *Chrysanthemum*, mango, and a combination of both ATSBs on the bacterial makeup of the mosquito midgut using hydrogel beads infused with these plants extract. According to our research, Proteobacteria and Firmicutes were the two most prevalent phylum in the midgut. Our finding states that *Ae. aegypti* and *Ae.albopictus* reared in the same laboratory harbor a different midgut bacterial microbiome due to the difference in the mosquito diets. By providing insights into the complex interplay between sugar sources and the mosquito microbiome, our research opens up new avenues for understanding the biology and disease transmission potential of these mosquitoes. Furthermore, our findings highlight the need for further research to confirm the significance of the observed bacteria and their potential impact on mosquito physiology and disease transmission.

## Acknowledgements

This research is supported by BioG Expert Sdn. Bhd. and Ministry of Higher Education Malaysia.

## Author Contributions

R.S: Investigation, experimental, data analysis and writing-original draft; N.F.A.K: Supervision, resources and project administration; W.M.W.A.K, S.S, K.Z.Z and W.R.W.I: Methodology, supervision, resources; C.E.W: Writing review and editing. All authors have read and agreed to the published version of the manuscript.

## Data Availability

All data included in this study is available upon request.

## Ethical statement

This study was approved by USM/IACUC/2022/(137)(1223).

